# *De novo* Genome Assemblies of Four Rainbow Trout Genetic Lines Reveal Structural Variants In Pursuit of a Pangenome Reference

**DOI:** 10.1101/2025.10.11.681701

**Authors:** Ali Ali, Geoffrey C. Waldbieser, Ramey C. Youngblood, Paul A. Wheeler, Brian E. Scheffler, Stuart Willis, Shawn R. Narum, Gary H. Thorgaard, Mohamed Salem, Yniv Palti

## Abstract

Rainbow trout (*Oncorhynchus mykiss*) exhibit extensive genomic diversity shaped by domestication, life history, and geographic origin. To advance the development of a comprehensive pangenome reference, we present new *de novo* genome assemblies of two genetically and ecologically distinct lines: Whale Rock (WR; wild, landlocked, Central California) and Keithley Creek (KC; wild, resident, interior Columbia Basin), along with the previously published assemblies of the Arlee (domesticated, Northern California) and Swanson (semi-domesticated, resident, Alaska) lines. All assemblies provide nearly complete coverage of known genes (BUSCO 95.8–99.7%) and are similar in genome size (∼2.3 Gb), with scaffold N50 values between 3.4 Mb (KC) and 52.4 Mb (Swanson). Comparative whole-genome alignments revealed high sequence conservation (97–98% identity) among assemblies, but also evidence of extensive structural variation of at least 50 bp in length. Structural variant (SV) profiling identified tens of thousands of deletions, insertions, and complex rearrangements largely in noncoding sequences. In an initial assessment of the utility of having multiple *de novo* genome assemblies for rainbow trout, we found that two strains (Arlee and Swanson; domesticated) share SVs enriched in genes linked with growth, reproduction, and adaptation to domestication, such as GTP binding and ECM-receptor interaction. In comparison, the other two strains (WR and KC; wild origin) share SVs associated with reproductive timing such as GnRH signaling pathway. Both Arlee and WR also have unique SVs potentially related to their geographic origin and unique life history. Additionally, we identified SVs in key regions, such as a QTL for fillet yield on Omy17 and the maturation-associated SIX6/ERβ-GPHB5 locus on Omy25q, suggesting the importance of considering SVs when investigating the genomics of complex traits. Together, these assemblies and comparative analyses establish a foundation for a rainbow trout pangenome reference, illuminating how they can be utilized to reveal the structural genomic basis of domestication, adaptation, and other complex traits in *O. mykiss*.

## Introduction

Rainbow trout (*Oncorhynchus mykiss*) originating from the west coast of North America have been introduced globally for aquaculture and recreational fishing [1]. It is one of the most widely cultivated fish species, where hatchery artificial selection dates back to the 1870s in California. While selective breeding for aquaculture and sport fishing have shaped the genetic makeup of domesticated stocks, anthropogenic and natural barriers (e.g., waterfalls, landslides, dams) have contributed to the geographic isolation of many of the wild populations. All these forces have contributed to the remarkable genomic and ecological diversity of the species [2–5].

As a highly adaptable salmonid, rainbow trout has significant ecological, economic, and evolutionary importance [6, 7]. The species has been widely used as a model for studying adaptation, domestication, and population genetics. Rainbow trout genetic lines have demonstrated differences in temperature tolerance [8], degree of domestication [9], behavior [10], development time [11], and disease resistance [12]. For example, genomic differences were found in IgL repertoire between lines of trout which could affect immune responses during disease [13]. Additionally, trout clonal lines exhibited significant differences in the relative expression of HSP70 under thermal stress, suggesting genetic differences underlying the variation [14]. Understanding the genomic basis of the variation is crucial for conservation efforts, sustainable aquaculture, and improving hatchery breeding programs.

Structural genome variation is broadly defined as insertion/deletions, inversions, duplications, and translocations that are larger than 50 bp [15]. While sequence-level (e.g. single-nucleotide polymorphism or SNP) variation has long been studied as a source of genetic differences between and within brood lines or populations, a growing body of research is demonstrating the large importance of structural variation in shaping important phenotypic variation in humans and in numerous domesticated and wild vertebrate species, including fishes [7, 16–18]. For example, a strong association between two adjacent inversions and migratory ecotypes was documented in Atlantic cod (*Gadus morhua*) [19]. Similarly, strong associations between inversions and adaptation to different habitats like freshwater and seawater were also found in other fish species like the threespine stickleback [20, 21]. Additionally, structural deletion variants were found to influence phenotypic variation in fish; for example, in sticklebacks, recurrent deletions of a *Pitx1* enhancer underlie pelvic reduction, highlighting how regulatory SVs can drive adaptive morphological changes [22]. Perhaps the most well documented association between structural genome variation and evolutionary adaptation in rainbow trout is the one between the large double inversion on chromosome 5 (Omy05) and sex dependent migratory behavior [7]. However, despite their large impact on genomic and genetic diversity in model organisms and the growing list of cases in wild fish populations, SVs are still understudied in most species, including rainbow trout. This is primarily due to the larger technical challenge and large investment in genome sequencing that is required for the detection and genotyping of SVs. To that end, multiple *de novo* genome assemblies and the generation of *pan-genome* references can provide invaluable resources for the detection of SVs in rainbow trout and for research on their association and impact on traits of interest. While originally applied to microbial species which exhibit a ‘core’ set of genes present across all individuals and a large library of variably present ‘accessory’ genes, the term *pan-genome* has been increasingly used in reference to plant and animals genomic resources as the unexpected degree of structural variation among individuals has become apparent. Herein, we use *pan-genome* to refer to a utilitarian compendium of the known small-scale (e.g. SNP) and large-scale (e.g. SV) variation among several to many individual genome assemblies of a species [23].

Recent advances in sequencing technologies have enabled the *de novo* assembly of high-quality reference genomes of non-model organisms. The rainbow trout genome assemblies from the Swanson and Arlee lines are notable examples that have been assembled using advanced sequencing methods [6, 7, 24–26]. This provided valuable insights into the species genomic architecture. The Swanson and Arlee genome assemblies have been used for studying SNPs, SVs, and other genomic features that contribute to phenotypic diversity and populations differentiation [24, 26–32]. For example, alignments of the chromosome sequences from the Swanson and the Arlee genome assemblies has revealed karyotype differences due to fissions of chromosome arms in the metacentric chromosomes Omy04, Omy14 and Omy25 and major inversions on chromosomes Omy05, Omy20, and Omy26 [6, 7, 26]. However, those two reference genomes do not fully represent the species’ genetic diversity which evolved as a result of multiple events leading to geographic isolation, domestication, and local adaptation [33, 34]. To fully capture the genomic landscape of rainbow trout, more efforts are still needed to assemble multiple reference genomes that will provide a more complete representation of the collective genetic diversity of the species.

In this study, we describe two new chromosome-level *de novo* genome assemblies for the Whale Rock (WR) and Keithley Creek (KC) genetic lines and present an initial examination of the breadth of genomic diversity across rainbow trout strains through comparative analysis of the four representative genomes. The strains from which those genetic lines originated have distinct ecological and evolutionary histories. Therefore, their genomes are expected to have unique genetic variants associated with adaptation to specific environments. The WR doubled haploid homozygous line originated from a wild landlocked steelhead population from the WR reservoir in coastal central California. The original anadromous steelhead population became landlocked by the dam construction that created the WR reservoir in 1961 and steelhead were recorded migrating from the tributaries into the reservoir as 1-and 2-year-olds in the 1970s and 1980s [35]. In addition, maturing adult fish from the WR line exhibit silver body coloration that is consistent with smoltification in steelhead (Thorgaard and Wheeler, Unpublished). Therefore, it is likely that the population from which the WR line was derived has retained migratory behavior and tolerance to marine or estuarine conditions, similar to the observed ability of landlocked steelhead descendants to produce viable smolts and adults after 70 years in freshwater [36]. On the other hand, the wild Redband trout from KC is native to inland river systems, and thus it represents the inland lineage of rainbow trout (*O. m. gairdneri*) in contrast to the three previous assemblies that were done using fish from the coastal lineage (*O. m. irideus*) [37]. By comparing these genomes with the previously assembled Swanson and Arlee strain genomes, we identify SVs, changes in gene content, and other genomic features that may underlie differences in their history of domestication, phylogeographic divergence, or local adaptation. Moreover, several recent studies have described genomic regions associated with life history variants important for population resilience in exploited or imperiled anadromous populations of *O. mykiss* (steelhead), including migration-timing, age-at-maturity, and repeat spawning phenology [38, 39]. However, these analyses relied exclusively on examinations of small-scale sequence variation (SNP and short insertion/deletions) based on whole genome data aligned to a single genome assembly. To further demonstrate the potential utility of a pan-genome reference for rainbow trout, we used comparative analysis of the four de-novo genome assemblies in those previously described genomic regions with the aim of identifying larger structural variations that are co-localized with the smaller sequence variants and that were otherwise undetected to date.

The overall aim of the comparative analyses we described in this manuscript was to demonstrate how *de novo* genome assemblies in rainbow trout can be used for the identification of conserved and strain-specific genomic regions, with the goal of providing the basis of a *pan-genome* for rainbow trout that can be used for future studies on the association between genome variants and phenotypes as well as for exploring the genetic basis of adaptation, domestication, and population structure.

## Materials and Methods

### Rainbow trout sampled for the genome assemblies

The Whale Rock (WR) rainbow trout we used for this *de novo* genome assembly was sampled from the established doubled haploid YY male line at Washington State University [27, 40]. Similar to the Arlee line, it has 2N=64 chromosomes and it originated from the Central California Coast. The one known major difference is that it originated from a wild fish from a landlocked steelhead population while the Arlee line was established from a domestic freshwater hatchery stock.

The Keithley Creek (KC) line was prepared using the established protocol to induce androgenesis with the aim of starting a doubled haploid line at Washington State University through irradiation of eggs obtained from an aquaculture stock and fertilization of the irradiated eggs with milt from a single wild KC rainbow trout male, which was followed by heat shock of 31.5C for 5 minutes that was applied 260 minutes post fertilization to block the first cell cleavage of the fertilized eggs [41]. However, since our DNA sequences mapping analysis in the process of the genome assembly indicated higher level of heterozygosity than in the established doubled haploid rainbow trout lines [40], we are obligated to caution that in the process that was used to establish the KC line at Washington State University, there might have been an inadvertent maternal contribution from the donor of the irradiated eggs, which was an aquaculture strain of rainbow trout possibly from a coastal origin.

### Genome sequencing

**WR:** DNA was extracted from a blood sample of a single WR YY male doubled haploid rainbow trout (NCBI Bio-sample Accession SAMN32933077) using methods as previously described [26]. Long reads DNA sequencing was performed using the continuous long reads (CLR) mode on the PacBio RS-II system (CD Genomics, Shirley, NY), generating 14.2 million reads with 254.7 Gb of DNA sequence data (106x coverage; NCBI SRA accession SRX19198672). An additional 5.5 million high fidelity (HiFi) reads with total length of 74.8 Gb (accessions SRX27459513, SRX27459514, SRX27459515, SRX27459516) were generated using circular consensus sequencing (CCS) mode on the PacBio Sequel II system at the USDA-ARS Genomics and Bioinformatics Research Unit (GBRU) in Stoneville, MS. In addition, Illumina sequence data was used for polishing and indel corrections of the assembly sequence scaffolds. A total of 558M whole genome Illumina paired-end reads (2×150bp; 70x coverage) were generated by a third party service provider (Admera Health, South Plainfield, NJ) using the Kapa Hyper Prep PCR free kit, and submitted to the NCBI SRA database (SRX21517362). The short-read sequences were processed with the program Trimmomatic (Version 0.38) to remove the Illumina sequencing adaptors.

**KC:** DNA was extracted from a blood sample of a single rainbow trout male fish from the KC line (NCBI Bio-sample Accession SAMN37796906). HiFi reads were produced using the CCS mode on the PacBio Sequel II system (GBRU Stoneville, MS). A total of 84.2 Gb sequence data (35x) was generated from 588,679 HiFi reads (NCBI SRA Accessions SRX22156623, SRX22156624, SRX22156625, SRX22156626).

### Assembly of contigs

Two different PacBio sequencing modes and contig assemblers were used for the two genomes. The WR line has a doubled haloid genome that is nearly 100% homozygous. Hence we followed the approach that we previously used for the genome assemblies of the Arlee [26] and Swanson [6] doubled haploid lines. This approach used the CLR PacBio mode that generates longer reads with higher base calling error rate and the Canu assembler to achieve an overall higher contiguity. For the KC contigs assembly we needed to generate two haplotypes since this genome was from a heterozygous diploid fish. Therefore, for the KC contigs assembly we chose the Hifiasm assembler with the HiFi reads from the CCS mode, which are shorter than the CLR mode reads but contain much lower error rate to enable haplotypes separation in heterozygous regions of the genome.

**WR**: PacBio long-reads were assembled into contigs using the Canu assembler (Version 2.2) [42]. The raw PacBio reads were aligned for error correction, and were trimmed of poor-quality segments and overlapping sequences, after which Canu retained 2.5 million high-quality reads with 84.6 Gb of DNA sequence data (35.3x). Canu assembled the retained reads into 1,479 contigs. The total length of the Canu contigs assembly was 2.34 Gb with a minimum, maximum, and N50 contigs length of 1.3 Kb, 53 Mb, and 10.4 Mb, respectively.

**KC**: The genome assembler Hifiasm [43] was used to generate two genome-wide haplotypes (Hap1 and Hap2) for the outbred diploid KC male rainbow trout using input data from the PacBio HiFi long read sequences and the HiC Illumina short reads. The total length of the Hap1 contigs assembly was 2.36 Gb in 6,120 contig sequences with a minimum, maximum, and N50 contigs length of 4.6 Kb, 10.4 Mb, and 1.1 Mb, respectively. The Hap2 contigs assembly was composed of 5,609 sequences for a total combined length of 2.32 Gb with a minimum, maximum, and N50 contigs length of 7.4 Kb, 12.1 Mb, and 1.1 Mb, respectively.

### Assembly of scaffolds

**WR:** The Bionano optical mapping with the Saphyr platform (Bionano Genomics, San Diego, CA, USA) was used to improve the contiguity of the assembly by generating the Bionano scaffolds assembly. DNA was extracted from a blood sample of the WR Male DH line fish and then labeled with Direct Label Enzyme (DLE-1). The Bionano Solve software (Solve3.3_10252018) (https://bionanogenomics.com/support-page/bionano-access-software/) was used to process the raw mapping data yielding a *de novo* Bionano assembly guided by the Canu contigs assembly. The Bionano hybrid scaffolds pipeline was then used to scaffold the Canu assembled contigs with the Bionano genome map, generating 1,215 scaffolds with a total length of 2.36Gb and N50 of 42Mb. We then used the PacBio HiFi sequence data to polish and correct sequence errors in the scaffold sequences. In the first step, we removed all the scaffolds that were smaller than 5Kb from the Bionano hybrid scaffolds assembly to retain 1,183 scaffolds. In the second step, the HiFi reads were mapped to the remaining scaffolds using pbmm2 (PacBio minimap2), and in the third step, we used Racon [44] to correct sequencing errors in the scaffolds assembly. The Illumina short-read data were mapped to the assembly scaffolds to correct indel errors following the methods we have previously described for the Arlee line genome assembly [26].

Chromosomal scale scaffolds were generated from the Bionano hybrid scaffolds assembly using Hi-C proximity ligation sequencing data. The Hi-C library was prepared from a frozen blood sample sequenced by a commercial vendor (Phase Genomics, Seattle, WA) to produce 780 million Illumina paired-end reads (2×150 bp) (SRA accessions SRX23202418 and SRX23202419). The Hi-C scaffolds assembly, including merging of contigs and scaffolds from the Bionano hybrid assembly and breaking of chimeric scaffolds, was then generated using the scaffolding program Salsa [45] following the same methods as we have previously described [26]. The Hi-C scaffolds assembly included 1,133 scaffolds with a total length of 2.35 Gb and N50 of 71.4 Mb.

**KC:** The Hi-C library was prepared from a frozen blood sample (Phase Genomics, Seattle, WA) to produce 762 million Illumina paired-end reads (2×150 bp) (SRA accessions SRX23159327 and SRX23159328). Hi-C scaffolding was prepared from the Hap1 contigs assembly that was a bit larger than Hap2 and was found to include the sdY gene and thus the Y chromosome. The contigs were checked with the NCBI fcs_adaptor program and 17 contigs were found to have the Biosciences Blunt Adapters. These adapters were trimmed at the end of nine contigs and in eight contigs they were removed by breaking the contig before and after the adapter. Here again the program Salsa was used to scaffold contigs together generating an assembly of 3,066 scaffolds with a total length of 2.36 Gb and N50 of 11.3 Mb.

### Assembly of chromosome sequences

**WR:** In order to produce chromosomal sequences from the Hi-C assembly scaffolds, linkage data from a SNP genetic map were used as previously described for the Omyk_1.0 [7] and OmykA_1.1 [26] chromosomal sequences assemblies. Briefly, the flanking sequences of each SNP marker were mapped to the scaffolds using the program NovoAlign (http://www.novocraft.com/products/novoalign/), which was then used to order, orient, and concatenate the scaffolds into 32 chromosomes using the genetic linkage data. Using the linkage information, eight large Hi-C scaffolds were broken into 19 smaller scaffolds. The final chromosome-level genome assembly (USDA_OmykWR_1.0; GCA_029834435.1) contains 102 scaffolds that were placed on 32 chromosomes and 1,036 unplaced smaller scaffolds, with a total sequence length of 2.3 Gb and N50 of 50.8 Mb. The karyotype of N=32 we identified for the WR male genome is consistent with the previously reported karyotype for the WR female doubled haploid line [46] and the coastal California origin of this genetic line.

**KC:** As described above for the WR male assembly, the genetic map linkage information was used to anchor, order, orient, and concatenate the scaffolds from the KC Hi-C assembly onto chromosome sequences. In the process, 90 Hi-C scaffolds that did not match the linkage map information were broken into 402 smaller scaffolds. The final chromosome-level genome assembly (USDA_OmykKC_1.0; GCA_034753235.1) contains 1,079 scaffolds that were placed on 30 chromosomes and 2,042 unplaced smaller scaffolds, with a total sequence length of 2.3 Gb and N50 of 3.4 Mb.

There is no specific information about the expected karyotype of the KC population. Some work that was done on other populations of the inland rainbow trout lineage (*O. m. gairdneri*) suggested a haploid karyotype of N=29 [47]. However, based on the number of major scaffolds and their unique mapping pattern to genetic linkage groups, and their sequence alignments with chromosome arms from the Arlee and Swanson genomes [6, 26], we concluded that the haplotype number of chromosomes for the individual fish that was sampled for this genome assembly is N=30. To that end, it is worth noting here that there might have been an inadvertent maternal contribution from an aquaculture strain of rainbow trout possibly from a coastal origin in the cross that was used to establish the KC line at Washington State University. Hence, we advise caution in drawing definitive conclusions regarding the karyotype and the haploid chromosome number of native rainbow trout from the inland lineage.

### Annotation of WR, KC, and Swanson assemblies

The WR, KC and new Swanson genome assemblies were annotated using Liftoff (v1.6.3) [48]. The Arlee genome assembly (USDA_OmykA_1.1) [26], and its corresponding gene annotation (GFF) file, was used as reference to lift genes from. Annotation parameters used were-s 0.9 (for minimum sequence identity), and-a 0.9 (for minimum alignment coverage). The-copies option was used to search for extra copies of genes that are not annotated in the reference after the initial lift over. The-chroms option was used to lift-over the annotations chromosome by chromosome, and then genes that did not map were aligned to the whole genome. A list of unplaced sequences was provided with the-unplaced option to map them to the target assembly after genes on the main chromosomes have been mapped. Unmapped features were then saved in a separate file with (-u). The-polish option was used to maximize alignments and improve annotation quality.

### Whole-genome alignment and SV detection using MUMmer and SyRI

To identify SVs (≥50 bp in length) among rainbow trout strains, we used MUMmer (v3.23) [49] and SyRI (v1.6.3) [50] for whole-genome alignment and variant calling. NUCmer (NUCleotide MUMmer) version 3.1 was applied for pairwise alignments of genomes using the -- maxmatch option under the following: minimum cluster length (-c) of 500 bp, break length (-b) of 500 bp, and minimum match length (-l) of 100 bp. Settings were selected based on prior studies [51, 52], and all values are reported in DNA base pairs (bp). Arlee was used as the primary reference genome in this study, except for three cases where WR genome served as a reference to identify wild-specific syntenic blocks, WR un-aligned/unique regions, and Arlee-unique SVs. For downstream analysis using DNAdiff (version 1.3), delta files were filtered using delta-filter with the following identity (-i 90) and minimum alignment length (-l 100) of 90% and 100 bp, respectively. Filtered delta files were used as input to dnadiff for alignment statistics and genomic differences. For calling SVs, delta files were subsequently filtered with-m option to retain 1-to-1 alignments, and coordinates were retrieved using show-coords with-THrd option. SyRI was then run with filtered delta and coordinate files using the reference and query FASTA sequences for the detection and annotation of SVs. All SVs included in downstream analyses were ≥50 bp in length. SyRI analysis commands were executed with at least two computational threads (--nc 2) and the logging configured for debugging (--log DEBUG). All SyRI analyses were conducted within Anaconda environment for reproducibility and for managing dependencies.

### SV annotation and filtering

For each line alignment with the Arlee or WR as the reference, chromosome-level VCF files that were generated by SyRI were compressed, indexed, and then concatenated with bcftools concat (v1.3.1). To functionally annotate SVs, we used an in-house Python pipeline to simultaneously annotate multiple VCF files using SnpEff v5.2. The script went through a specified directory for indexed VCF files. For each input VCF, the pipeline produced an annotated VCF as well as complementary HTML and CSV summary reports. Annotations were performed using a custom SnpEff database derived from the *O. mykiss* Arlee reference genome assembly (GCF_013265735.2) [26] and corresponding GTF annotation file. SnpEff annotated the SVs as MODIFIER (non-coding/untranslated regions or RNA gene) or functional with LOW (minor changes), MODERATE (changes in coding sequence), or HIGH (severe changes including stop/start gains and feature truncations) effects. To identify SVs with potential functional consequences, we used SnpSift (version 5.2) to filter high-impact insertions, deletions, inversions, and duplications across the three query genomes. The six VCF files with filtered SVs for each line based on alignment to the Arlee and WR genome as the reference were deposited in Dryad (datadryad.org) and will be publicly released after peer review and acceptance of the manuscript.

### Gene enrichment analysis

To investigate the biological functions associated with genes affected by SVs, gene enrichment analysis was performed using ShinyGO v0.77 (https://bioinformatics.sdstate.edu/go77/) [53]. Lists of unique gene identifiers affected by a specific type or category of SVs were uploaded to ShinyGO, setting *O. mykiss* as the target species. Default parameters were used, including a false discovery rate (FDR) cutoff of 0.05 for assessing statistical significance. The software generated interactive plots and tables of enriched pathways and terms, utilized to understand the potential biological impacts/consequences of SVs of interest.

### Large-effect regions for *O. mykiss* life history variants

Several analyses were conducted to examine the degree to which structural variation may be identified in genomic regions that were previously found to be associated with life history variations in *O. mykiss*. First, we made alignments of the large-effect regions of the four assemblies, which reside in chromosomes Omy28 (migration-timing) and the q-arm of Omy25 (age-at-maturity and repeat spawning phenology), using Mummer v4 (nucmer --nosimplify-c 100-g 1000) to examine the presence of structural differences in these regions. Second, for the Omy25 region, we selected 16 stocks (populations) for which low coverage whole genomic data was available and which represent the diversity of steelhead stocks within the Columbia River Basin [54]. We mapped this sequence data to each of the four assemblies using *bwa* v0.7.18 [55], filtered discordant and improperly paired alignments using *samtools* [56, 57], and examined the mean coverage across the adjacent regions in Omy25q associated with age-at-maturity and repeat spawning phenology, respectively. Within these regions, we calculated in R (R Corp.), from mpileup files of each stock produced by *samtools,* the mean coverage in 500bp sequential windows, and then normalized this coverage to the mean coverage for each stock across the entire region.

## Results and Discussion

### *De novo* genome assemblies

The *de novo* genome assemblies analyzed in this study were derived from four genetic lines of *O. mykiss* (rainbow trout). The four lines exhibited variation in chromosome numbers, geographic origin, domestication, and life history (Table 1) making them valuable resources for comparative genomics and the study of complex traits in *O. mykiss*. For example, the variation in chromosome numbers (ranging from 58 to 64) highlights the diversity of karyotypes that was captured by the selection of those four lines as the foundation towards developing a pan-genome reference panel for rainbow trout.

**Table 1.** Genetic lines used for the *de novo* genome assemblies.

| Line Name | 2N* | Sex | Geographic Origin | Life History Type | Origin |
| --- | --- | --- | --- | --- | --- |
| WR Male | 64 | YY Male | Central California Coast | Landlocked Steelhead | Wild |
| Arlee | 64 | YY Male | Northern California | Resident | Domesticated |
| Swanson | 58 | YY Male | Kenai Peninsula, Alaska | Resident | Semi-Domesticated** |
| KC | 60 | XY Male | Columbia Basin, Snake River | Resident | Wild |

\* A diploid chromosome number (2N); \*\*Fish have undergone partial domestication after one generation in the hatchery environment.

The assembly statistics of the four rainbow trout genomes revealed notable similarities and differences in assembly quality and continuity (Table 2). All four assemblies have the same genome size (2.3 Gb), ungapped length (2.3 Gb), GC content (43.5%), and chromosome level sequences that were guided using the robust genetic linkage information. This indicates consistency in the overall structure and completeness of the four assemblies. However, the KC genome assembly is the most fragmented, likely due to the lower genome coverage (35.0x), the use of a heterozygous individual fish that is not a doubled haploid, and the application of shorter PacBio HiFi reads instead of high-coverage PacBio CLR sequencing to generate the initial contigs level assembly. The KC assembly has significantly more gaps (1,049), scaffolds (3,121), and contigs (6,120), as well as the lowest scaffold N50 (3.4 Mb) and contig N50 (1.1 Mb) values. The Swanson and WR are the most contiguous assemblies. Both assemblies have high scaffold N50 values (52.4 Mb and 50.8 Mb, respectively) and low L50 values (16), indicating fewer and larger scaffolds. The Arlee assembly also exhibited high continuity, with the fewest scaffolds (938) and contigs (1,228), as well as the highest contig N50 (15.6 Mb). Both the Swanson and WR genome assemblies were enhanced by the PacBio CLR and HiFi reads, supplemented by multi-platform scaffolding using Bionano optical mapping and Hi-C proximity ligation information. While Swanson and WR have the best overall assembly quality, Arlee shows strong contiguity, and KC is the most fragmented of the four assemblies. Together, these results suggest that sequencing depth, read quality, and the combination of multiple sequencing and scaffolding technologies all contributed to the quality differences between the KC and the other three rainbow trout genome assemblies.

**Table 2.** Genome assemblies’ statistics.

| **Feature** | **OmykA_1.1**<br>**(GCA_013265735.3)**<br>**(Arlee)** | **OmykWR_1.0**<br>**(GCA_029834435.1)**<br>**(WR)** | **Omyk_2.0**<br>**(GCA_025558465.1)**<br>**(Swanson)** | **OmykKC_1.0**<br>**(GCA_034753235.1)**<br>**(KC)** |
| --- | --- | --- | --- | --- |
| Number of chromosomes | 32 | 32 | 29 | 30 |
| Total length | 2.3 Gb | 2.3 Gb | 2.3 Gb | 2.3 Gb |
| Number of scaffolds | 938 | 1,138 | 1,278 | 3,121 |
| Gaps in scaffolds | 486 | 382 | 460 | 2,999 |
| Gaps between scaffolds | 196 | 109 | 70 | 1,049 |
| Scaffold N50 | 39.2 Mb | 50.8 Mb | 52.4 Mb | 3.4 Mb |
| Scaffold L50 | 22 | 16 | 16 | 182 |
| Number of contigs | 1,228 | 1,520 | 1,738 | 6,120 |
| Contig N50 | 15.6 Mb | 12.2 Mb | 10.7 Mb | 1.1 Mb |
| Contig L50 | 41 | 55 | 57 | 541 |
| Genome coverage | 111.6x | 94.3x | 106.0x | 35.0x |
| BUSCO score* | C: 99.7%<br>(S: 55.5%; D: 44.2%) | C:96.2%<br>(S:47.2%,D:49.0%) | C:96.5%<br>(S:48.0%,D:48.5%) | C: 95.8%<br>(S:47.2%,D:48.6%) |

\*Benchmarking Universal Single-Copy Orthologs (BUSCO) using version 3.1.0 and lineage actinopterygii\_odb9. C: Percent of complete protein-coding genes detected. S: Percent in Single copy. D: Percent of Duplicated genes.

### Intronic and intergenic SVs dominate the landscape across the rainbow trout lines

To identify SVs between the lines we aligned the genome assemblies of KC, Swanson, and WR to the Arlee reference genome (Additional File 1: Table S1). The alignment statistics revealed notable differences in the variants landscape between the lines. All three lines showed high sequence conservation with the reference genome. The average identity values range from 97% to 98%. However, SV analysis revealed a wide range of genomic rearrangements. Overall, the Swanson genome assembly exhibited the highest number of indels, breakpoints, and relocations, as well as frequent translocations and insertions relative to the Arlee genome. In contrast, KC and WR are more conserved relative to the Arlee genome, exhibiting fewer relocations and genomic variations such as inversions. These differences suggest that the Swanson line may represent a more genetically distinct population, but it is important to note here that the difference in number of variants between Swanson and the other three genomes is inflated by the fission of Omy04, Omy14, and Omy25, as well as the large double inversion on Omy05, for which Swanson is the only genome with the Rearranged (R) haplotype, while the other three genomes exhibit the Ancestral (A) haplotype.

SyRI annotated a range of SVs, including inversions, translocations, duplications, unaligned regions, insertions, and deletions. SVs were filtered based on size thresholds using a custom Python script (Figure 1a). Insertions, deletions, and tandem repeats were retained if ≥50 bp [15], whereas other complex SVs were filtered using a more stringent threshold of 100 bp to reduce false positive calls [58–60]. Notably, deletions and insertions were the most abundant SV types across all comparisons. The counts ranged from 49,533 to 55,334 for deletions and 48,310 to 53,807 for insertions. Such high counts reflect wide sequence variation and mobile element activity between the three query strains and the reference genome [61, 62]. Additionally, high duplications and un-aligned region frequencies were detected. This might be an indication of potential lineage-specific expansions or reference collapse [63, 64]. Complete list of SV counts in the three rainbow trout lines are provided in Table S2 (Additional File 1). As a positive control to validate the SV detection pipeline, we aligned the X chromosome (Omy29) from the recently assembled hybrid-derived haploid genome of a rainbow trout from the domesticated Hayspur hatchery strain (GCA_049308825.1) [65] to the Y chromosome of the Arlee reference genome. The MUMmer/SyRI-based approach successfully identified a 169,949 bp deletion encompassing the sdY gene, which is known to be absent in XX rainbow trout females [66].

**Figure 1.**
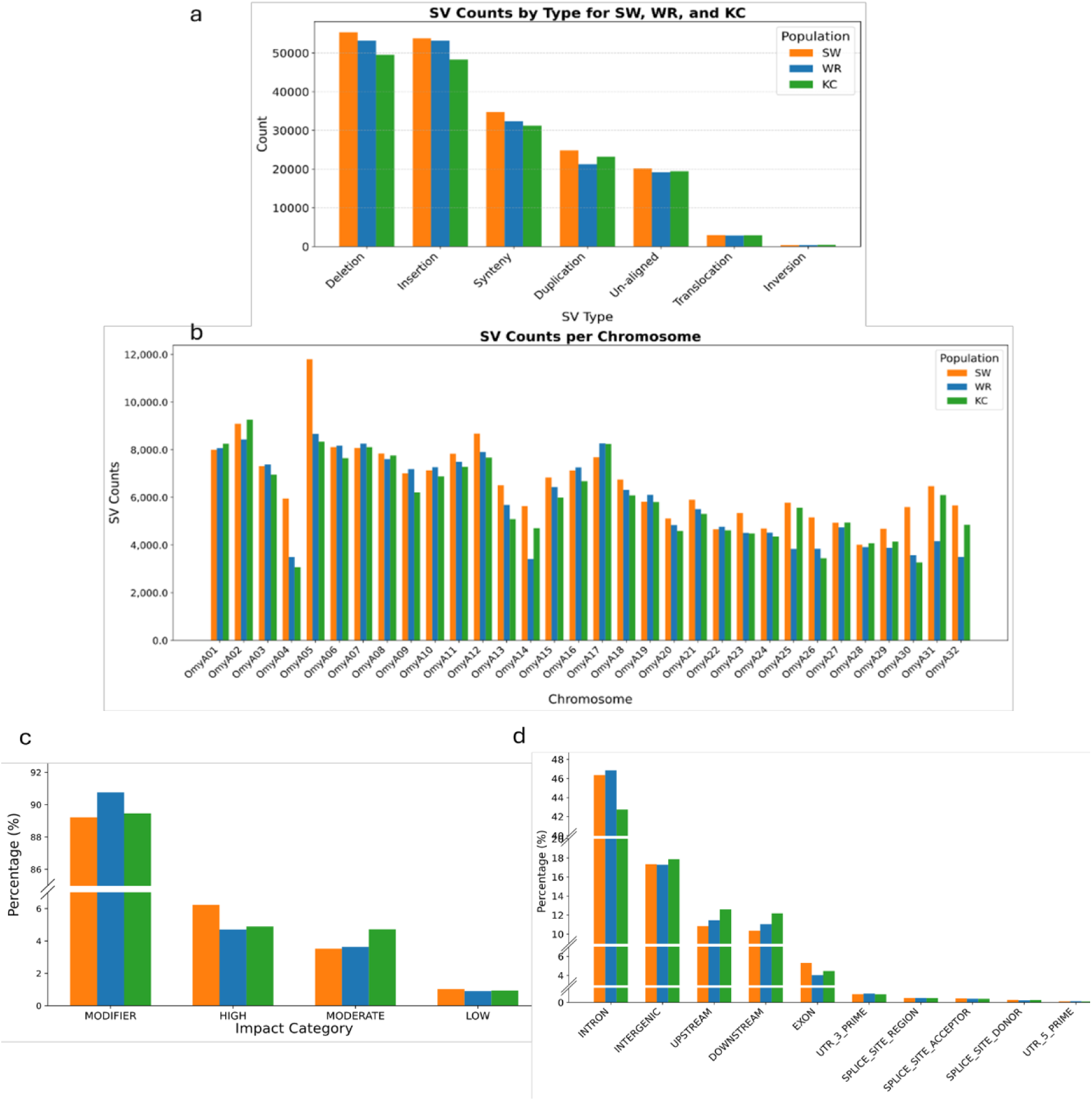
Characterization of SVs identified in the Swanson, WR, and KC lines in reference to the Arlee genome assembly. a) Distribution of filtered SV types annotated by SyRI in the reference-query alignments reveals a predominance of insertions and deletions. b) Total SVs counts per chromosome. c) Annotated SV frequencies in each effect class, which were dominated by the MODIFIER class. d) Genomic features affected by SVs, indicating that the majority of variants are in intronic and intergenic regions with a smaller fraction impacting coding and regulatory regions.

Functional annotation of the SVs revealed consistent patterns in variant type, impact, and genomic distribution among the three non-reference rainbow trout lines—Swanson, WR, and KC (Figure 1b-d). The three lines showed a consistent pattern in variant distribution across chromosomes. Population-specific differences in variant counts were observed on Omy05 and chromosomes that underwent fission (Omy04, Omy14, and Omy25). Swanson and WR tend to show slightly elevated variant counts on some chromosomes compared to KC (Figure 1b). Most of the variants were from the MODIFIER impact category (>89%) and localized in non-coding regions without predicted functional consequence. LOW-and MODERATE-impact variants occurred in a minor fraction (∼0.9–3.5%), and HIGH-impact variants were even more infrequent (∼4.7–6.2%) (Figure 1c). Genomic feature analysis identified that SVs largely affected intronic (∼42.8–46.9%) and intergenic regions (∼17%), and fewer being in exons, untranslated regions (UTRs), or splice sites (Figure 1d). Collectively, these findings demonstrate a highly conserved SV landscape across the three lines, with most variants likely to have low functional effects but with a subset potentially having regulatory or coding effects of interest. These results are similar to those of Liu et al. (2021) [29] that conducted SVs discovery and characterization in rainbow trout using short reads whole genome sequencing and found that 95% of the SVs in their study were classified in the MODIFIER category and genomic locations identified predominantly as intronic (45.5%) or intergenic (25.3%) based on alignments to the previous reference genome assembly (Omyk_1.0) [7]. Furthermore, in a previous study we also showed that intron retention is the most frequent alternative splicing event in rainbow trout [24], which suggests that SVs are more likely to influence gene regulation or splicing rather than disrupting or modifying coding regions directly in this species.

Further analysis of high-impact SVs revealed differences in SV characteristics and burden among the three rainbow trout lines when aligned to the Arlee reference genome (Table 3). A total of 724 high-impact SVs were identified in KC, compared to 657 in WR and 598 in Swanson. Inversions were less frequent whereas duplications were the predominant SV type across the three lines (> 90%). As expected, Swanson displayed the largest SV (35,109,550 bp), a large double inversion, located on Omy05. Chromosomal hotspots for SV accumulation differed among populations: OmyA17 in KC, OmyA01 in WR, and OmyA13 in Swanson. A substantial number of unique genes were affected, with KC showing the highest count (4,982). Functional impact analysis indicated that duplications, gene fusions (including bidirectional fusions), frameshift variants, and stop-gained mutations with duplications were common effect types.

**Table 3:**
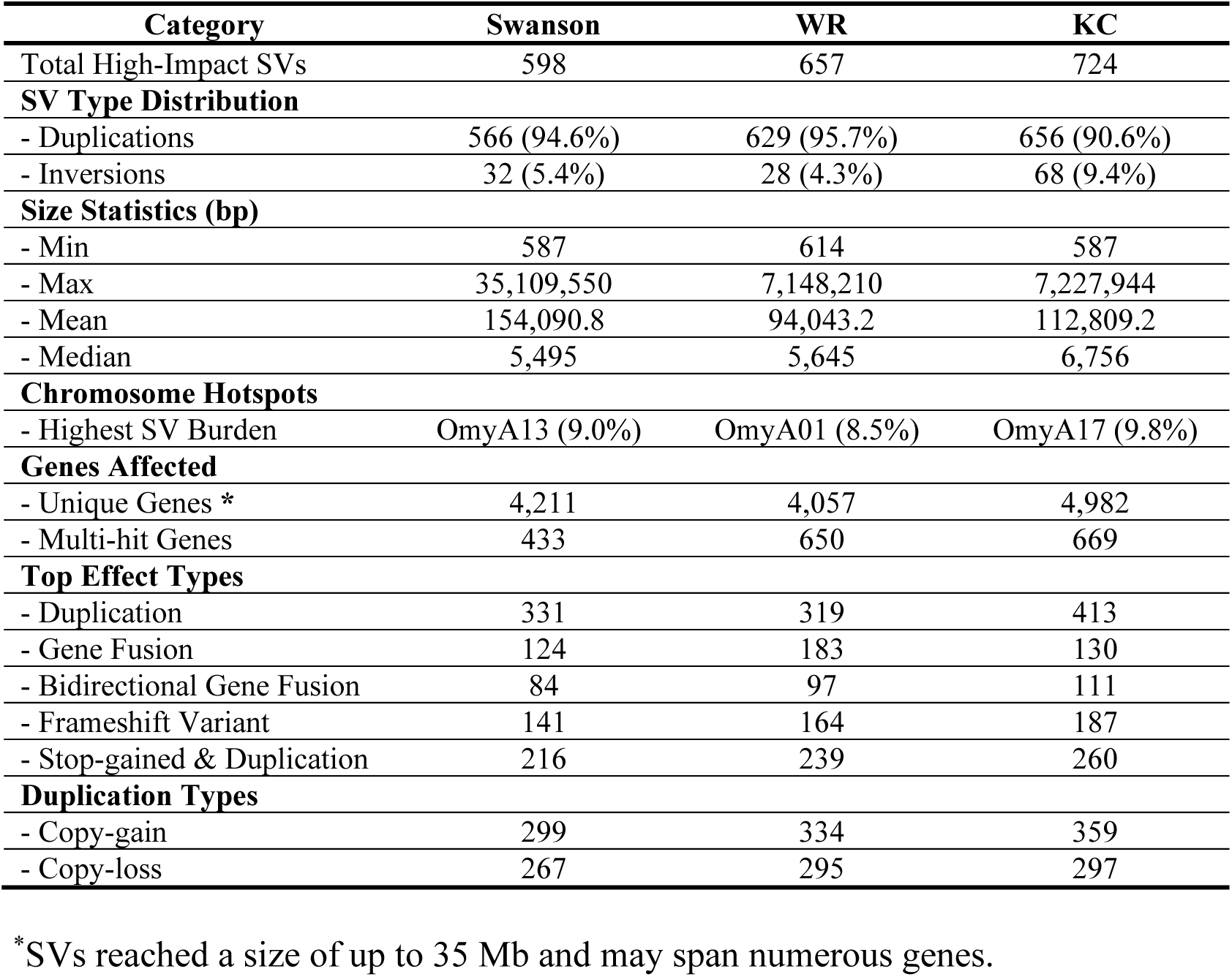
Summary of high-impact SVs identified in KC, WR, and Swanson compared to the Arlee reference genome.

### Comparative SVs profiles between the rainbow trout lines

To further investigate the biological relevance of SVs among strains, we grouped SVs into their presence or absence within domesticated and wild lineages, as well as between the WR line that represents a landlocked steelhead population and the other three lines that represent freshwater resident strains of origin. This grouping enabled a preliminary identification of SVs for strains that differ by (1) domestication history and (2) life history background. Four general groups (Table 4) were formed:

**Table 4:**
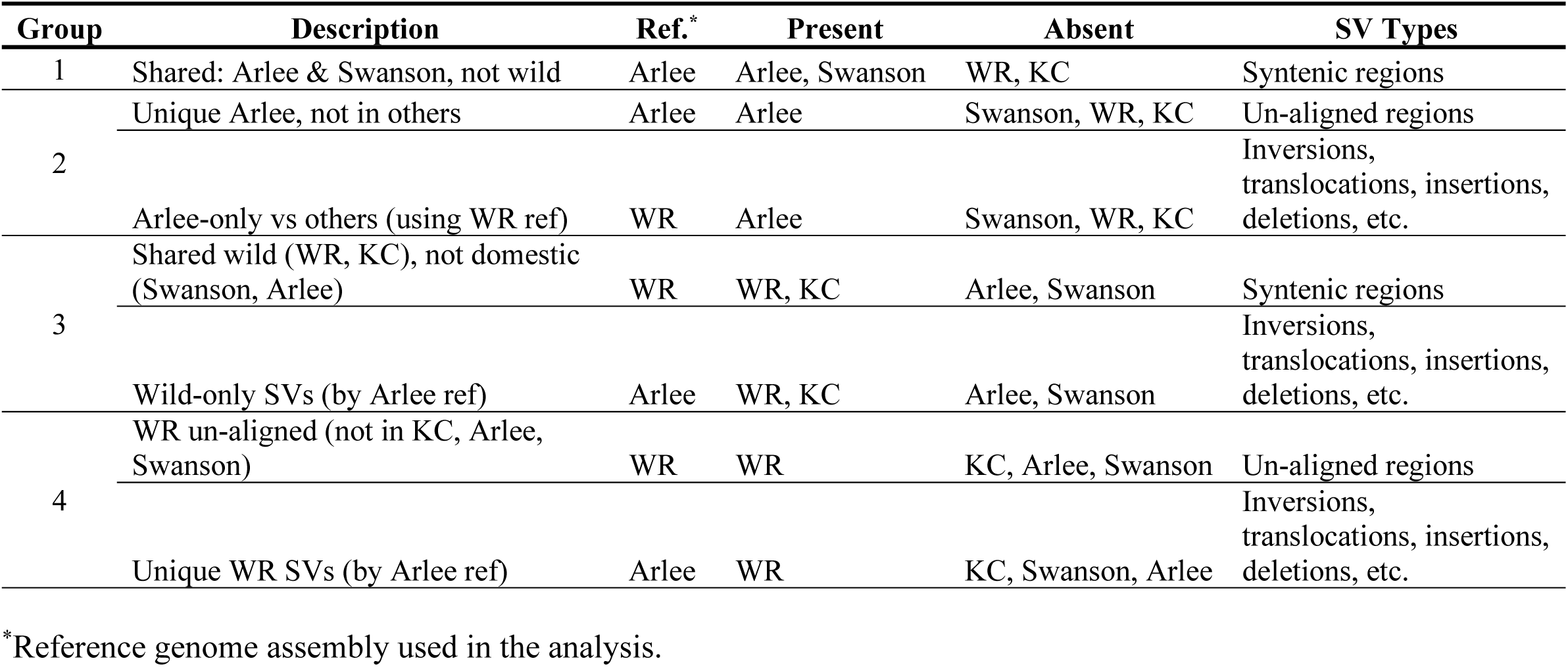
Classification of SVs by strain of origin or shared life history.

1. SVs present in Arlee and Swanson but not in the lines derived from wild strains (WR and KC).

This group had a combined total of 404 syntenic blocks shared by Arlee and Swanson but not by WR and KC (Table S3). These SVs had a maximum length of 175,589 bp (Average size 21,205 bp). Notably, chromosome OmyA13 had the highest proportion of these SVs, at 27.5%, which reflects a region of genomic divergence among strains.

A total of 289 syntenic regions (∼72%) shared only by Arlee and Swanson overlapped 339 genes, among which the coding sequences (CDS) of 245 genes were completely encompassed by 195 of these syntenic regions. Functional annotation of the 245 genes revealed several enriched pathways (Table S4). Interestingly, enrichment for nucleotide binding functions, including GTP binding, ribonucleotide guanyl binding, and guanine nucleotide binding were detected. Guanylate binding proteins (GBPs) are involved in cell proliferation and have been associated with growth traits in cattle [67], which may suggest a potential role in selective breeding practices in captivity. Additionally, variations in G protein-coupled receptors have been linked to domestication traits, including reproduction, development, and stress responses [68]. However, further examination with larger sample sizes from domesticated vs wild strains will be necessary before inferences can be made regarding domestication effects.

2. Arlee-unique SVs not found in other strains may represent strain-fixed SVs resulting from artificial selection, genetic drift, or potential introgression.

We found 4,269 genes annotated in 7,298 Arlee-specific SV regions unmapped to Swanson, WR, or KC (i.e., un-aligned regions) (Table S5). Of them, the CDS of 2,095 genes were encompassed by the un-aligned regions (> 95% coverage). The highest proportion of the Arlee-unique blocks was on chromosome OmyA13 (7.0%). Notably, two genes located within un-aligned regions—LOC118940322 (*CD22*) on Omy17 and LOC118943595 (*TAAR6*) on Omy22— matched candidate gene presence/absence variations (PAVs) previously identified in American-selected rainbow trout [69]. *TAAR6* (Trace Amine-Associated Receptor 6) is related to the G protein-coupled receptor family that was found enriched in group 1. However, gene set enrichment analysis of all Arlee-specific genes within un-aligned regions (Table S6) identified the significant overrepresentation of the ECM-receptor interaction pathway (FDR = 0.0417; Fold Enrichment = 6.26), which contains genes, such as *itgb3b*, *SV2A*, and *lamc3*. The ECM-receptor interaction pathway has been linked to reproductive traits in chickens, which affect the efficiency of spermatogonial stem cell development and quality of meat [70, 71]. Efficiency in reproduction and meat quality are preferred traits in domesticated animals, and hence ECM pathways may play a role in these traits.

Additionally, we detected 169,030 distinct Arlee line SVs (inversions, translocations, duplications, insertions, deletions, and other complex SVs) using the WR as the reference for genome alignment. The average length of the SVs is approximately 1,691 bp, with the largest variant at 4,089,394 bp. The CDS of 41 genes associated with fatty acid synthesis, energy metabolism, and enzymatic activity were fully encompassed within the inserted loci. Similarly, the CDS of 10 other genes were completely overlapping deleted loci in Arlee. Among them, the *Semaphorin receptor*, *Plexin-A1*, which regulates axon guidance, perhaps due to less need for complex environmental navigation. No enriched pathways were identified for genes in which less than 95% of their coding sequences were affected by indels.

Several key biological pathways were recurrently enriched across diverse SV categories. This included insertions (Figure 2a), deletions (Figure 2b), and also duplications, inversions, translocations, and other complex SVs. Of note, Focal adhesion, MAPK signaling, Neuroactive ligand–receptor interaction, Endocytosis, and Adrenergic signaling pathways were each associated with at least seven different SV types (Table S7). The Focal Adhesion pathway (FDR = 2.36E-41) was significantly enriched, involving 131 genes in cell adhesion and signal transduction. This is consistent with findings in avian models, where elevated activity in the liver of line S fowl under artificial selection for growth was mainly involved in focal adhesion, ECM-receptor interaction, cell adhesion molecules, and signal transduction, collectively contributing to enhanced muscle development and lipid metabolism in Guangxi Partridge chickens [72]. In addition, we noticed enrichment of pathways with potential impacts on energy metabolism in the heart and cardiac function like the Adrenergic signaling in cardiomyocytes (FDR = 3.24E-25). The presence of overlapping genes such as *AKT1*, *PIK3R1*, *RAF1*, *SRC*, and *GRB2* across multiple pathways underscores their central regulatory roles. These pathways combined collectively indicate participation in vascular development, cell communication, growth, and stress responses [73–81]. Overall, this comparative analysis of the *de novo* genome assemblies demonstrates how SVs with a potentially profound impact on phenotypic diversity, may be associated with the processes of local adaptation or artificial selection.

**Figure 2.**
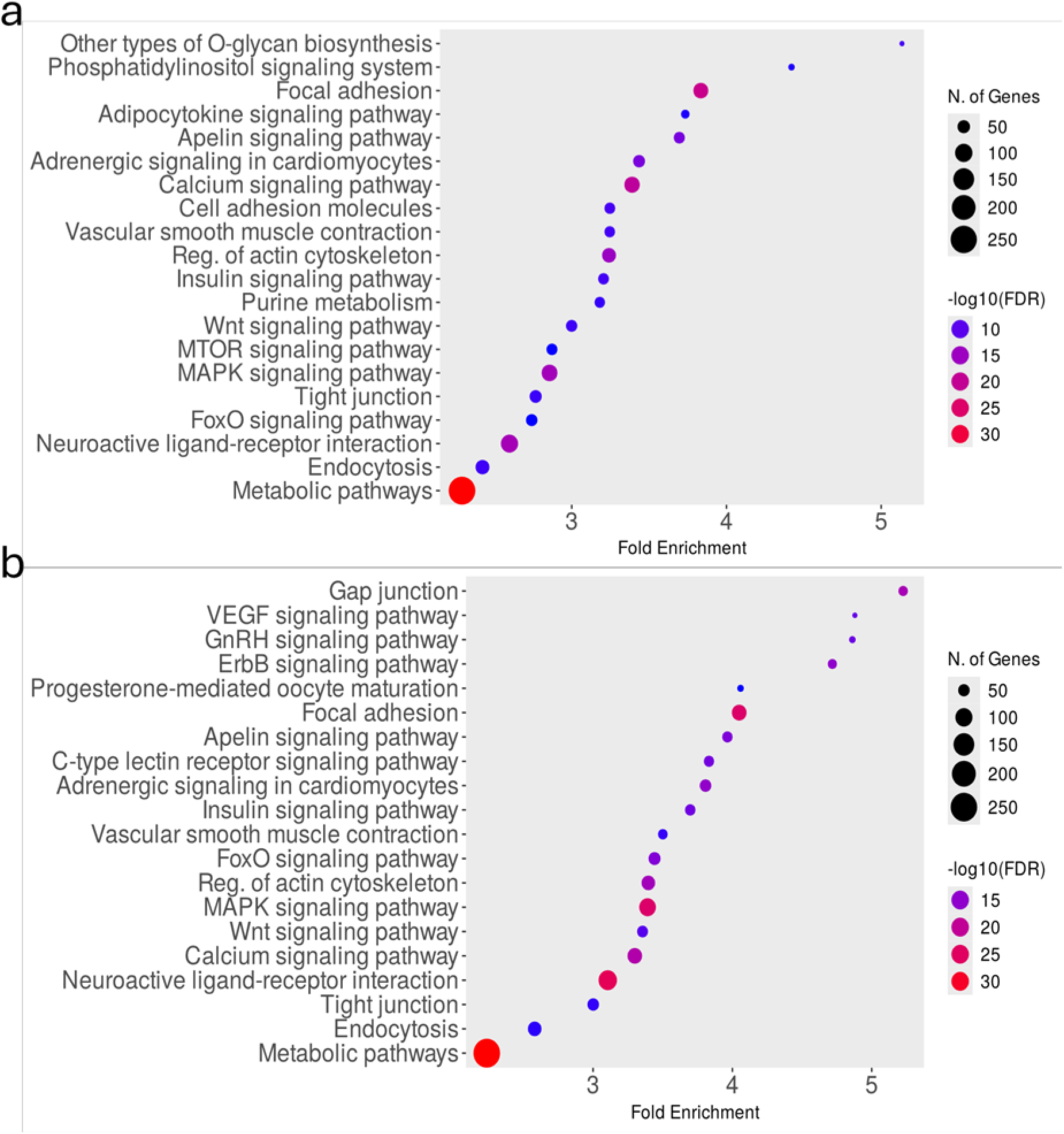
Gene enrichment analyses of genes affected by insertions (a) and deletions (b) unique to the Arlee line.

3. SVs present in WR and KC, absent in Arlee and Swanson.

We compared syntenic regions that are shared by the wild strains (KC and WR) but not in the domesticated lines (Arlee and Swanson) (Table S8). There were 62 such regions ranging from 880 bp to 55,501 bp (median: 6,660.5 bp; mean: 8,779.8 bp). Chromosome OmyA17 had the highest proportion of such wild-specific syntenic regions that accounted for 32.2% of the total. A total of 28 syntenic regions shared only by wild strains overlapped 110 genes, among which the CDS of 17 genes were completely encompassed by 18 of these syntenic regions. Functional annotation revealed several pathways, including Adherens junction, Tight junction, Focal adhesion, and Regulation of actin cytoskeleton (Figure 3a).

**Figure 3.**
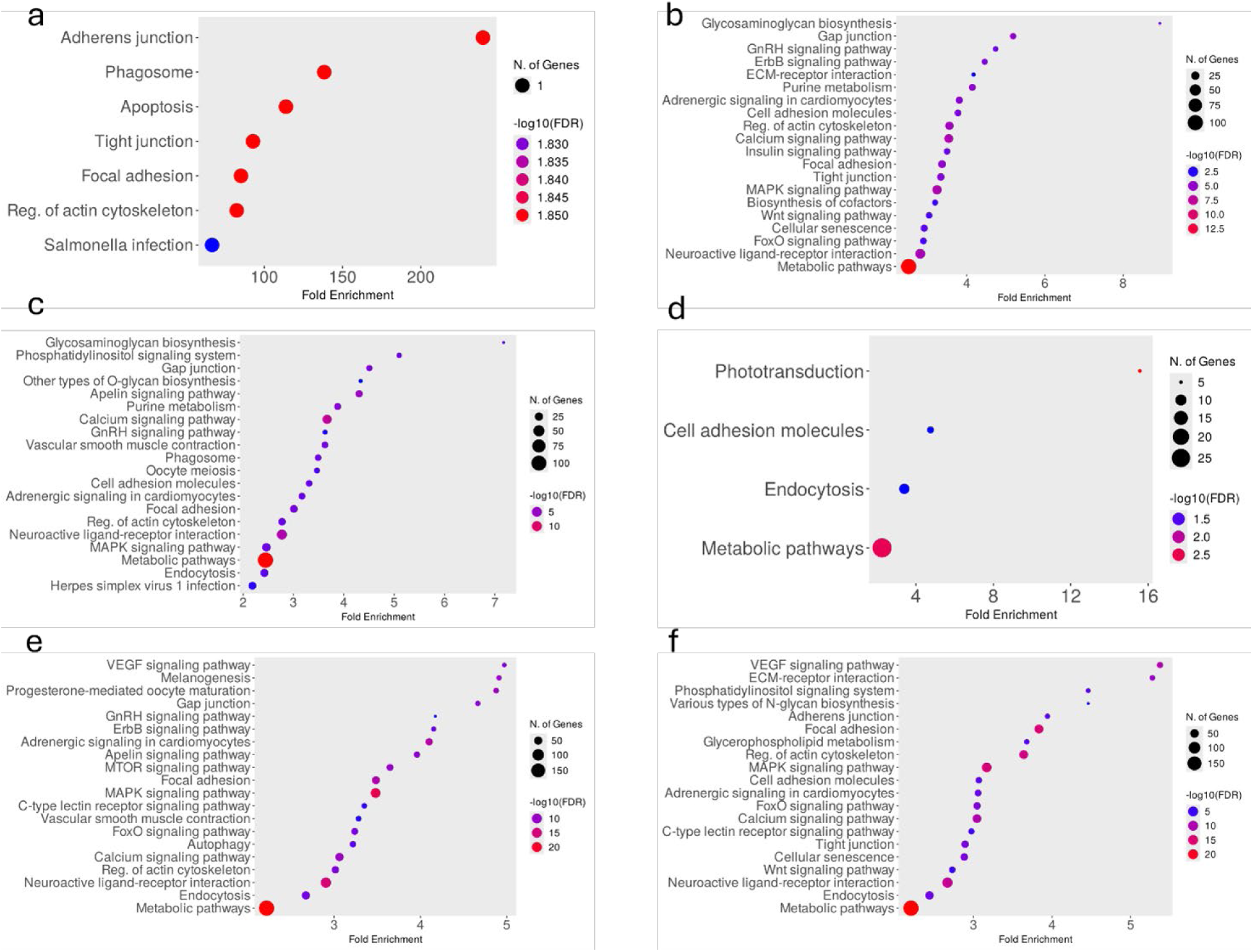
Gene enrichment analyses of genes encompassed or affected by syntenic regions (a), insertions (b), and deletions (c) unique to WR and KC, the lines with ‘wild’ strains of origin, and absent or not affected in Swanson and Arlee, the lines derived from the more domesticated strains. d-f) Gene enrichment analyses of genes encompassed or affected by un-aligned regions (d), insertions (e), and deletions (f) unique only to the WR strain.

Other than un-aligned and syntenic regions, we detected 56,479 SVs (deletions, insertions, inversions, translocations, etc.) shared between wild strains WR and KC but not in the domesticated strains Arlee and Swanson. The average size of these SVs is approximately 2,629 bp and the longest form covers 3,652,591 bp, showing a broad variation in SV sizes. These wild-restricted SVs may reflect the limited diversity represented by only four fish or natural variation that has been lost to domestication or evolution to the wild environment.

Similar to group 2, pathways related to environmental responsiveness, neurosensory processing, and stress adaptation (Table S9) were recurrently enriched across diverse SV categories such as insertions (Figure 3b), deletions (Figure 3c), duplications, and other complex SVs. These pathways include adrenergic signaling in cardiomyocytes, Calcium signaling pathway, Gap junction, Metabolic pathways, MAPK signaling pathway, Neuroactive ligand-receptor interaction. Notably, MAPK signaling pathway (FDR = 1.14E-19) and Calcium signaling pathway (FDR = 1.53E-16) are highly enriched, both of which are key to transducing extracellular environmental signals into appropriate cellular responses, such as those required for thermal tolerance, osmoregulation, and immune defense [82–86]. The significant enrichment of the Neuroactive ligand-receptor interaction pathway (FDR = 3.80E-15) and Adrenergic signaling in cardiomyocytes (FDR = 8.63E-12) suggests enhanced neurosensory and cardiovascular regulation [87–91] for predator evasion, foraging, and migration in natural populations.

Insertions encompassed 3,234 genes, but only the CDS of 66 genes were affected. Similarly, deletions spanned 3,606 genes of which CDS of 98 genes were impacted. This suggests that noncoding regions are impacted by indels. Interestingly, genes associated with the GnRH signaling pathway, vital for reproductive timing, were enriched in both insertion and deletion categories, suggesting potential evolutionary divergence in developmental and physiological regulation among the lines represented. Collectively, these findings demonstrate how the comparative analysis of *de novo* genome assemblies may provide insights on differences in the genomic organization between populations of rainbow trout.

4. WR–specific SVs.

To explore further genomic features potentially linked to life history of WR, we found structural regions in the WR genome that were un-aligned to the resident strains (KC, Arlee, and Swanson) (Table S10). There were 4,850 unique un-aligned regions with sizes up to 539,026 bp (median: 2,136 bp; mean: 14,763.9 bp). Chromosome 17 has the highest percentage of these regions (6.6%). WR-specific sequences could be genomic components that are associated with life history traits involved in local adaptation, or lineage-specific structural divergence, or merely a reflection of limited representation of genomic variation due to the small sample size used in this study. Further characterization of the 2,894 genes located within these genomic intervals revealed significant enrichment in pathways such as phototransduction, cell adhesion molecules, and endocytosis (Figure 3d). These findings demonstrate how comparative analysis of *de novo* genome assemblies may help uncover mechanisms underlying ancestral migratory behavior or other attributes resulting from the unique ecological pressures to which the steelhead populations are exposed.

Additionally, we identified 126,411 WR line unique SVs (inversions, translocations, duplications, insertions, deletions, and other complex SVs). These SVs have a maximum size of 1,224,483 bp, indicating that there are some extremely large structural alterations that potentially would have significant impacts on gene function or regulation.

There were a number of signaling pathways recurrently showing significant enrichment among the SVs from the WR line (Figure 3e-f; Table S11). MAPK signaling pathway, VEGF signaling, and Focal adhesion pathways ranked among the top enriched pathways, suggesting important functions in stress response, vascular remodeling, and cell adhesion. These processes are essential for physiological transformation during freshwater–saltwater transitions and long-distance migration [92–97]. The MAPK signaling pathway (FDR = 3.79E-28), which comprises over 120 genes such as *MAPK14*, *RAF1*, *AKT1*, and *NFATC1*, is responsible for cell response to environmental stresses like osmotic stress and temperature changes during migration [98–102]. Similarly, VEGF signaling pathway (FDR = 4.07E-20), including *vegfaa*, *src*, and *pik3r1*, is crucial for angiogenesis and vascular permeability [103, 104] which are important for sustaining elevated metabolic demands and oxygen delivery during periods of high activity [105, 106].

Focal adhesion (FDR = 3.79E-31) was also highly enriched. *PXN*, *ITGA6*, and *RAC1* genes from this pathway can be suspected of involvement in muscle plasticity and tissue remodeling, which are critical for swimming movement [107–110]. Regulation of actin cytoskeleton and Apelin signaling pathway suggests additional importance of cardiovascular functionality, and contractility of muscles, and osmoregulatory control [111–113]. Genes such as *mylk3*, *pdgfra*, and *pak6b* may reflect the physiological acclimatization of anadromous steelhead to fluctuating salinity environments. Activation of the ErbB, Insulin, and FoxO pathways also indicates neuroendocrine and metabolic regulation of migratory behavior [114–116]. These pathways regulate energy homeostasis, cell growth, and survival and are in agreement with known migratory traits such as enhanced feeding before seaward migration and increased stress resistance. Finally, Adrenergic signaling and Gap junction pathways (*adrb1*, *adcy5*, and *GJD2*) point to cardiac function, communication between neurons, and arousal behavior, each of which is essential for initiating and sustaining migratory behavior [117–119]. Collectively, the enriched pathways may represent preliminary evidence for local adaptation or divergent lineages, but we cannot exclude that they may merely be the result of limited representation of genomic variation in this sample.

Together, SVs observed among these four lines may represent drivers of genomic and phenotypic diversity across highly divergent lineages of rainbow trout. Several enriched pathways such as MAPK signaling, neuroactive ligand-receptor interaction, and calcium signaling are common among all comparison groups. This might occur when different SVs affect different genes within the same pathway, particularly in large pathways which involve many genes such as Neuroactive ligand-receptor interaction (488 genes) and MAPK signaling pathway (366 genes). Such overlap suggests the versatility/adaptability of conserved pathways to potentially mediate artificial or natural selection, where SV architecture and regulatory divergence may account for lineage-specific adaptation. The findings are in agreement with studies showing that salmonid SVs have a tendency to repurpose ancient pathways for novel ecological or domestication-related traits [18, 120].

### Exploring the overlap between identified SVs and QTLs for Muscle yield

We previously identified a quantitative trait locus (QTL) on Omy17, in which 50-SNP genomic windows accounted for ≥1% of additive genetic variance for fillet yield [28]. Notably, alignment of the KC and WR genome assemblies to the Arlee reference identified a chromosomal inversion on Omy17, not found in the Swanson assembly (Figure 4a-d). The inversion interval fully overlaps the QTL associated with variation in fillet yield in a domesticated aquaculture population. This suggests a potential functional link between the inversion and phenotypic variation observed. Similarly, previous studies reported inversions on Omy05 and Omy20 associated with genetic divergence between different ecotypes in rainbow trout [121, 122]. Gene annotation in the inverted region (OmyA17: 36,299,758 – 43,447,967) uncovered several biologically relevant candidates (Table S12), including *CMP-N-acetylneuraminate-beta-galactosamide-alpha-2,3-sialyltransferase 2* (glycosylation), *cytochrome c oxidase subunit 6C-1* (mitochondrial respiration), and *phosphoenolpyruvate carboxykinase 1* (a gluconeogenesis critical enzyme). In addition, the region has genes for cellular stress response pathways (*reactive oxygen species modulator 1*), proteasomal degradation (*E3 ubiquitin-protein ligase Itchy*), and transcriptional regulation (*NFX1-type zinc finger-containing protein 1* and *CBFA2T2*). The presence of *1,25-dihydroxyvitamin D(3) 24-hydroxylase*, a vitamin D metabolism gene, also indicates the potential metabolic relevance of this inversion.

**Figure 4.**
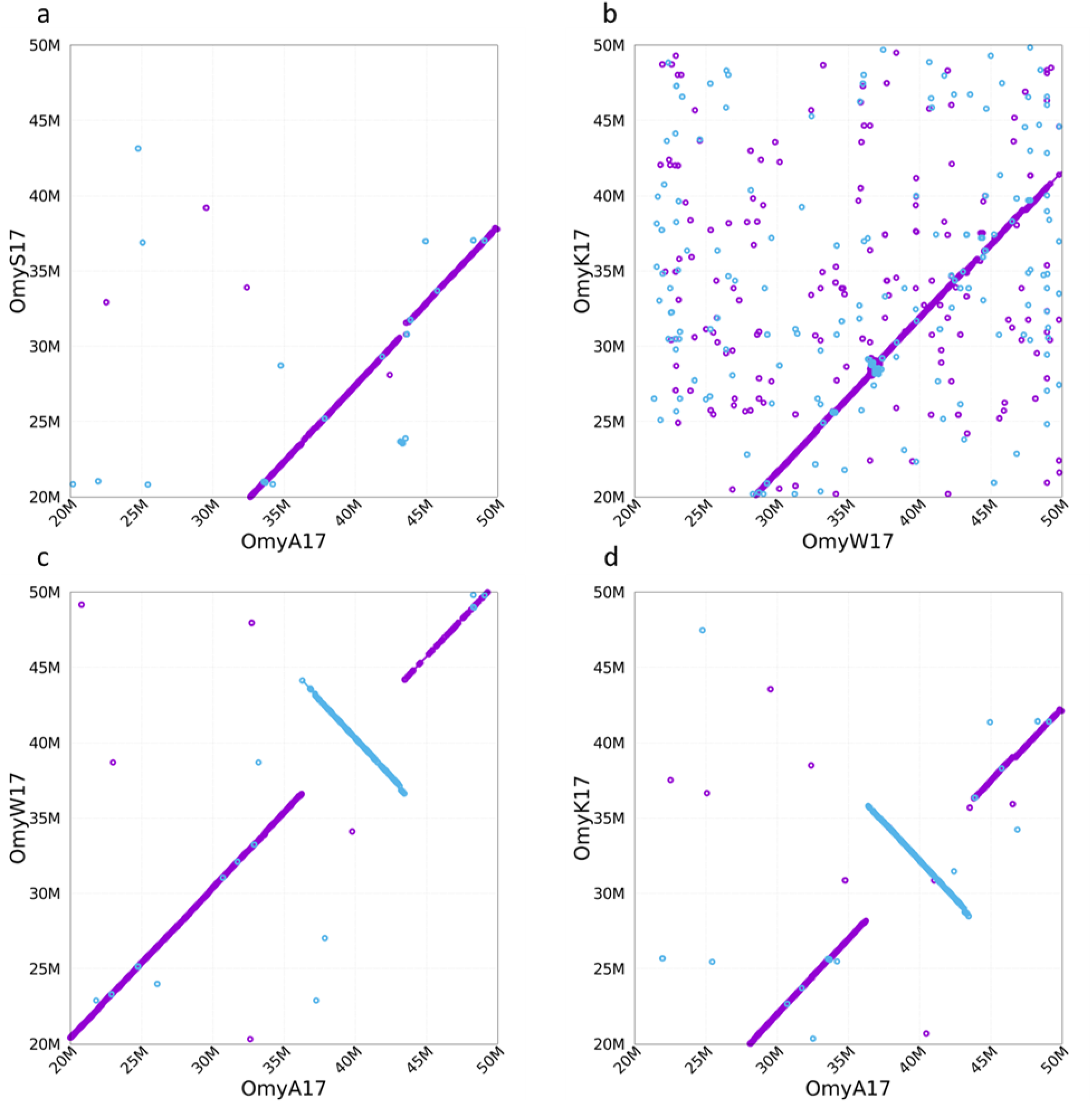
a) OmyS17 of the current Swanson assembly (Omyk_2.0) aligned to Arlee assembly OmyA17 and showing no inversion in the Omy17 QTL region. b) OmyK17 and OmyW17 of wild strains also showing no inversion. c) OmyW17 and d) OmyK17 aligned to the Arlee OmyA17 and showing an approximately 7 Mb inversion overlapping with the muscle yield QTL region.

Notably, the inversion is found in the wild KC and WR strains but not the fully domesticated Arlee and semi-domesticated Swanson lines. These findings suggest the Omy17 inversion might be a candidate SV underpinning physiological or other traits that warrant further investigation with a larger sample size of domesticated and wild rainbow trout populations to establish the significance of the potential association of this SV with domestication or with artificial selection in rainbow trout.

### Exploring structural variation in large effect loci for life history traits

Previous genome-wide association analyses for salmonids, including rainbow trout, have typically identified genomic regions with significant effect on important life history traits, using small variants (SNPs and small INDELs) inferred from a single genome assembly. Here, we looked for structural variation (SV) in two regions previously associated with migration timing and age-at-maturity/spawning phenology on chromosomes Omy28 and Omy25, respectively, in rainbow trout. Though we did not explicitly test for association of SVs with these traits, we discovered potentially important variation that would go unrecognized and untested in a typical GWAS analysis.

Examining alignments among the four assemblies for the large effect regions, we observed notable differences between the alignments (Additional File 2: Figure S1). In the Omy28 region, which includes the GREB1L and ROCK1 genes, there were some INDEL discontinuities between the assemblies, and which lay in the intergenic region between the two genes or within the introns of GREB1L. While these regions do not obviously cause major changes to the function of either of these genes, association studies (based on SNPs in previous genome assemblies) have shown that the intergenic and upstream (center-adjacent) portion of the GREB1L gene show the strongest association with migration timing in Columbia Basin steelhead [38], suggesting the intergenic structural variation could be similarly associated with this trait.

The region of Omy25q presents an even more complex case than Omy28. Willis et al. [39] found that two aspects of spawning in steelhead, age-at-maturity (age at maiden spawning) and repeat spawning phenology (proclivity to skip or spawn consecutively following maiden spawning), were separately associated with adjacent regions on Omy25q that occupy a contiguous island of linkage disequilibrium in two wild populations. Similarly, Willis et al. [54] noted that these regions exhibited exceptional allele frequency variation among 74 population samples from the Columbia Basin (Additional File 2: Figure S2). Alignments of this region among the four *de novo* genome assemblies showed more complex structural variation than the Omy28 region (Figure 5, Additional File 2: Figures S3-S6). In the region near the SIX6 gene, associated with age-at-maturity, structural variation (INDELs) was evident in the upstream intergenic region, which was also the region of highest association in the SNP-based GWAS analyses (Figure 5, Additional File 2: Figures S3-S6). Notably, this was also reflected in plots of relative windowed-mean coverage as differing areas of low or zero coverage across samples. While there was some coverage variation evident among population samples, it was small relative to the differences among genome assemblies (Figure 5, Additional File 2: Figure S7).

**Figure 5.**
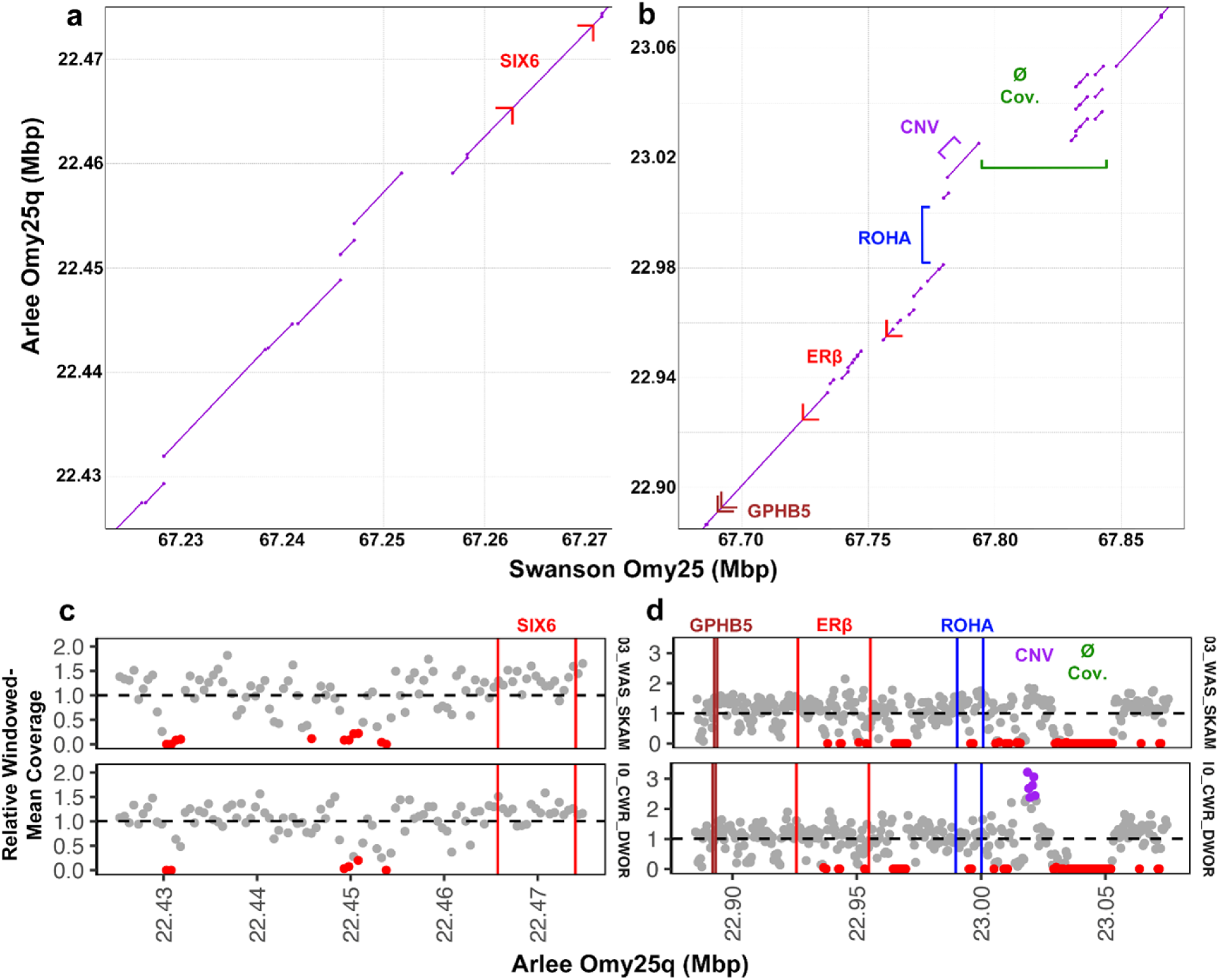
Structural variation in the chromosome Omy25q regions association with age-at-maturity (left) and repeat spawning morphology (right). a,b) Mummer alignments of subsets of the Omy25 chromosome from the Swanson and Arlee genome assemblies, with structural features labeled (genes are shown with right angles indicated the direction of transcription; ØCOV: zero read coverage; CNV: copy number variant; ROHA: region of highest association). c,d) Plots of relative windowed-mean sequence coverage for two Columbia Basin stocks on the Arlee genome assembly, with structural features labeled (Skamania Hatchery, Washougal River; Dworshak Hatchery, Clearwater River). Note that for (a,c) the entire region reflects the ROHA. The figure depicts structural genome variations detected from the alignment of the Arlee genome assembly to the Swanson assembly, but similar structural variations were detected in the alignment of the Arlee assembly to the other two de novo genome assemblies (Supplemental Figure S3).

Alignments of the four assemblies near the Omy25q region associated with repeat spawning phenology showed very complex variation upstream of two genes, ERβ and GPHB5, including both INDELs and duplications (Figure 5, Additional File 2: Figure S8), as well as a prominent INDEL in the first intron of the ERβ gene. Notably, the region most strongly associated with this trait is only present in the Arlee assembly and absent from the other three assemblies. Similarly, plots of relative windowed-mean coverage across populations revealed not only that the positions of zero coverage differed among assemblies, but also that there exists an adjacent copy-number variant (CNV) that varies in frequency among populations. In contrast, while plots of SNP frequency across this region show differences among stocks (Additional File 2: Figure S9), they do not capture the differences in structural variation, which may show a different pattern from SNP variants in mediating the adjacent genes and associated traits.

## Conclusions

This study provides a comparative analysis of *de novo* genome assemblies of four genetically diverse rainbow trout lines, an important foundational step toward the construction of a robust pangenome reference for *O. mykiss*. Our results show pervasive SVs including deletions, insertions, and complex rearrangements with lineage-specific patterns. Specifically, the genome regions that harbor the SVs were found to overlap with genes that are functionally enriched for growth, reproduction, and environmental adaptation pathways, implicating their potential roles in genetic diversity across divergent lines of this species. SVs were identified in key regions such as the Omy17 muscle yield QTL and the maturation-associated SIX6/ERβ-GPHB5 region, which highlights the role that they may have in the genetics of complex traits in rainbow trout. Our findings demonstrate the importance of high-quality, strain-specific genome assemblies that will provide comprehensive representation of intra-specific genetic diversity. This work paves the way for follow-up functional validation and evolutionary studies, and broadens the applicability of a pangenome framework for basic and applied research in salmonid genomics by highlighting the need to incorporate additional *de novo* genome assemblies from strains shaped by distinct life histories or other traits of interest.

## Acknowledgments

We thank Dr. Guangtu Gao with our utmost gratitude. Sadly, Dr. Gao passed away last year. Before his passing, he co-managed the sequence data collection and archiving and generated the *de-novo* genome assemblies of the WR and KC lines. We would like to take this opportunity to acknowledge his incredible talents and most impactful contributions to this work and to the past decade of salmonids genomics research. We thank Roseanna Long from USDA-ARS in Leetown, WV for helping with processing the DNA samples and Sheron Simpson from USDA-ARS in Stoneville, MS for technical assistance with the PacBio sequencing of the KC DNA sample. This study was supported by funding from Agricultural Research Service (ARS) in-house project number 8082-10600-002-000D. Mention of trade names or commercial products in this publication is solely for the purpose of providing specific information and does not imply recommendation or endorsement by the U.S. government. USDA is an equal opportunity provider and employer.

## Author Contributions

YP and GG conceived and planned the study. GT conceived the idea of generating doubled haploid lines that represent the species diversity for genetic and genomic analyses and supervised the collection, generation and propagation of the lines. PW was involved in gametes collection, and generated and propagated the lines. GT, PW, and SN provided the genetic lines and samples for this study. AA planned and conducted the SV analyses, and drafted the manuscript including tables, figures and supplemental information. YP planned, coordinated and supervised the DNA sequencing and data collection, handling and storage of the data, genome assemblies and analyses of the data, and contributed to the draft manuscript writing. GW, RY and BS conducted optical mapping and DNA sequencing and contributed resources. SW and SN conducted data analyses and contributed to the manuscript writing and revising and to the presentation of the results. MS provided resources and supervision, contributed to data interpretation, and revising the manuscript. All authors read and approved the final manuscript.

## Data Accessibility Statement

All the Whale Rock line raw genome sequence data and the final genome assembly are associated with the NCBI SRA BioProject Accession: PRJNA928430. All the Keithley Creek line genome sequence data and the genome assembly are associated with the NCBI SRA BioProject Accession: PRJNA1027447. The genome assembly accessions are: USDA_OmykA_1.1: GCA_013265735.3; Omyk_2.0: GCA_025558465.1; USDA_OmykWR_1.0: GCA_029834435.1; USDA_OmykKC_1.0: GCA_034753235.1.

The six VCF files with filtered SVs for each line based on alignment to the Arlee and WR genome as the reference were deposited in Dryad (datadryad.org) and will be publicly released after peer review and acceptance of the manuscript.

All the data analysis results are provided in the supplemental data files.

## Benefit-Sharing Statement

Benefits from this research accrue from the sharing of our data and results on public databases as described above.

## Additional File 1: List of Supplemental Tables

**Table S1:** Whole-genome alignment statistics of Swanson, Keithley Creek, and Whale Rock assemblies against the Arlee reference genome.

**Table S2:** Table S2: Counts of SyRI-annotated structural variants in rainbow trout strains before and after size-based filtration

**Table S3:** Syntenic regions present in domesticated strains (Arlee and Swanson) but absent in wild (Keithley Creek and Whale Rock).

**Table S4:** Functional enrichment analysis of genes with CDS completely encompassed by syntenic blocks shared by Arlee and Swanson but absent in wild strains (Whale Rock and Keithley Creek).

**Table S5:** Un-aligned genomic regions specific to Arlee not shared with Keithley Creek, Whale Rock, or Swanson.

**Table S6:** Gene enrichment analysis of genes annotated in Arlee-unique structural regions not aligned to Swanson, Keithley Creek, or Whale Rock.

**Table S7:** Functional enrichment analysis of genes located within regions of structural rearrangements unique to the Arlee strain and absent in Swanson, Keithley Creek, and Whale Rock.

**Table S8:** Syntenic regions present in wild (Keithley Creek and Whale Rock) but absent in domesticated strains (Arlee and Swanson).

**Table S9:** Functional enrichment analysis of genes located within regions of structural rearrangements shared by wild strains (Whale Rock and Keithley Creek) but absent in Arlee and Swanson.

**Table S10:** Un-aligned genomic regions specific to Whale Rock not shared with Keithley Creek, Arlee, or Swanson.

**Table S11:** Functional enrichment analysis of genes located within regions of structural rearrangements specific to the Whale Rock and absent in Swanson, Arlee, and Keithley Creek.

**Table S12:** Genes located within the Omy17 inversion interval overlapping the fillet yield QTL.

## Additional File 2: List of Supplemental Figures

**Figure S1.** Alignment of the GREB1L-ROCK1 region of Chr. 28 across assemblies.

**Figure S2.** Allele frequency differences on Chr. 25q among 74 populations of steelhead. (Modified from Willis et al. 2023).

**Figure S3.** Alignment of the SIX6-ERb region of Chr. 25q across assemblies.

**Figure S4.** Annotation and structural variation of the SIX6 and ERb region of Chr. 25q in alignment of the Swanson (2.0) and Arlee assemblies.

**Figure S5.** Annotation and structural variation of the SIX6 and ERb region of Chr. 25q in alignment of the Arlee and Whale Rock assemblies.

**Figure S6.** Annotation and structural variation of the SIX6 and ERb region of Chr. 25q in alignment of the Arlee and KC assemblies.

**Figure S7.** Relative sequence coverage by population across the 4 assemblies. Regions of low coverage are shown in red.

**Figure S8.** Relative sequence coverage by population across the 4 assemblies. Regions of low coverage are shown in red.

**Figure S9.** Allele frequencies across stocks at the ERB-GPDH5 region for each Omy assembly.

